# Barrier-free liquid condensates of nanocatalysts as effective concentrators of catalysis

**DOI:** 10.1101/2022.02.10.480015

**Authors:** Silky Bedi, Gaurav Kumar, S M Rose, Sabyasachi Rakshit, Sharmistha Sinha

## Abstract

Molecular confinement of catalysts enhances the catalytic activity significantly. However, physicochemical barriers in traditional confinements restrict the free-passage of substrates/products. To obtain a barrier-free confinement of catalysts, here we explored the liquid-liquid phase separation. Using favourable ionic strength and crowding agents, we recruit the protein-inorganic-composites in phase separated liquid condensates from a solution. The phase separation propensity of these nanocomposites was seen to be independent of the native conformation of the component protein. Using standard catalytic oxidation-reduction reactions, we show that the close-proximity yet diffusive nature of catalysts in solution amplifies the homogeneous catalytic-efficiency of metal particles significantly. Overall, our work demonstrates the roadmap of using inorganic catalysts in homogeneous homogenous solution phase with amplified efficiency and longevity.

## Introduction

Once the limit in catalytic efficiency is achieved from the variations in shape and size of catalytic materials in the nano-regime ^[1]^, interests to amplify the efficiency further are switched towards the spatiotemporal confinement of nano-catalysts ^[2]^. Microenvironments in the form of molecular vessels or crucibles of different architectures like core-shell structures, porous materials, 2-dimensional materials, and organic casings have been developed that demonstrated enhanced catalytic properties ^[3–6]^. Furthermore, caging helps to isolate nano-catalysts in a solution. However, physical barriers in cages often restrain the widespread applications of such trapped nano-catalysts, especially for bulkier substrates, and in solution-phase. A majority of catalytic applications is, therefore, still limited to heterogeneous catalysis on a solid substrate. Recently membrane-less condensates formed by process of liquid-liquid phase separation (LLPS) have been reported to enhance the enzymatic rate of reaction. LLPS is a spontaneous, de-mixing of a solution in two liquid phases, a condensed phase and a dilute phase. The two phases originate from the preferential, weak and spatially distributed multivalent non-covalent interactions among solutes in a solution leading to non-stoichiometric assemblies of solutes in the form of droplets ^[7]^. In cells, such LLPS of proteins is considered as the common origin of the membraneless compartments/organelles like nucleoli^[8]^, Cajal bodies^[9]^, stress-granules^[10]^ etc. The phenomenon of LLPS not only explains the formation of subcellular organelles but may also hold the potential to develop condensed membraneless phases with confined enzymes ^[11]^. As individual bio-reactors, these liquid droplets compartmentalize and confine biological reactions, and demonstrate significantly enhanced enzymatic turnover. Recruitment of enzyme and substrate in phase separated scaffold increases the SUMOylation reaction by 36-folds ^[12]^.

We hypothesized an enhanced catalysis of inorganic catalysts if they can be isolated and condensed to membraneless liquid droplets from a solution. In the present work, we set out to form liquid droplets of protein-coated inorganic nano-materials by tuning the microenvironment of the solution. Protein used to coat the nanomaterials would respond to change in the surrounding, facilitating the phase separation of protein-metal nanocomposite. We have used two structurally different proteins, namely α-synuclein and bovine serum albumin (BSA) to induce the formation of liquid droplets. α-synuclein is an intrinsically disordered protein and is known to phase separate in solution ^[13]^. BSA is a structured globular protein that has been used in several paradigms as crowding agents for phase separation ^[14]^. Using a combination of microscopic and spectroscopic studies we identify the appropriate ionic strength and macromolecular crowding of the surrounding solution that trigger the LLPS of these protein-based nanomaterials. We assessed differences in the catalytic proficiency of the synthesized nanomaterial in bulk and phase separated environments. Through our results, we bring out the ability of phase separated nanocatalysts to outperform the ones present in bulk environments.

## Results

### Liquid-liquid phase separation of BSA-Au nanocomposites

We begin our experiments by testing the condition to induce liquid-liquid phase separation in nanocomposites coated with BSA (**Fig 1a)**. BSA serves as a crowding agent during LLPS events in biology. In the initial experiments we used BSA coated gold nanoclusters [BSA@AuNCs] and BSA coated gold nanoparticles [BSA@AuNPs]. A broad absorbance spectra is observed in the UV-visible range for the BSA@AuNCs **(Fig.S1a)** indicating a nanocluster size of less than 3 nm ^[15]^. The BSA@AuNCs shows emission maxima at 720 nm when excited at 450 nm **(Fig.S1a)**. In contrast, the absorption spectra of BSA@AuNPs shows a plasmonic band at 520 nm **(Fig.S1b)** indicating bigger size. The CD spectra of the composites show lower ellipticity as compared to native BSA, suggesting alteration in the native structure of the protein upon conjugation with AuNCs and AuNPs **(Fig.S1c)**. Further, we noticed reduced intensities at 1650 cm^-1^ in FTIR for BSA@AuNCs and BSA@AuNPs compared to BSA alone **(Fig.S1d).** 1650 cm^-1^ in FTIR corresponds to α-helix content in protein. TEM imaging of BSA@AuNCs shows its size is in the range of 3.0 ± 0.5 nm as represented in **(Fig. S2)**.

**Figure 1:**
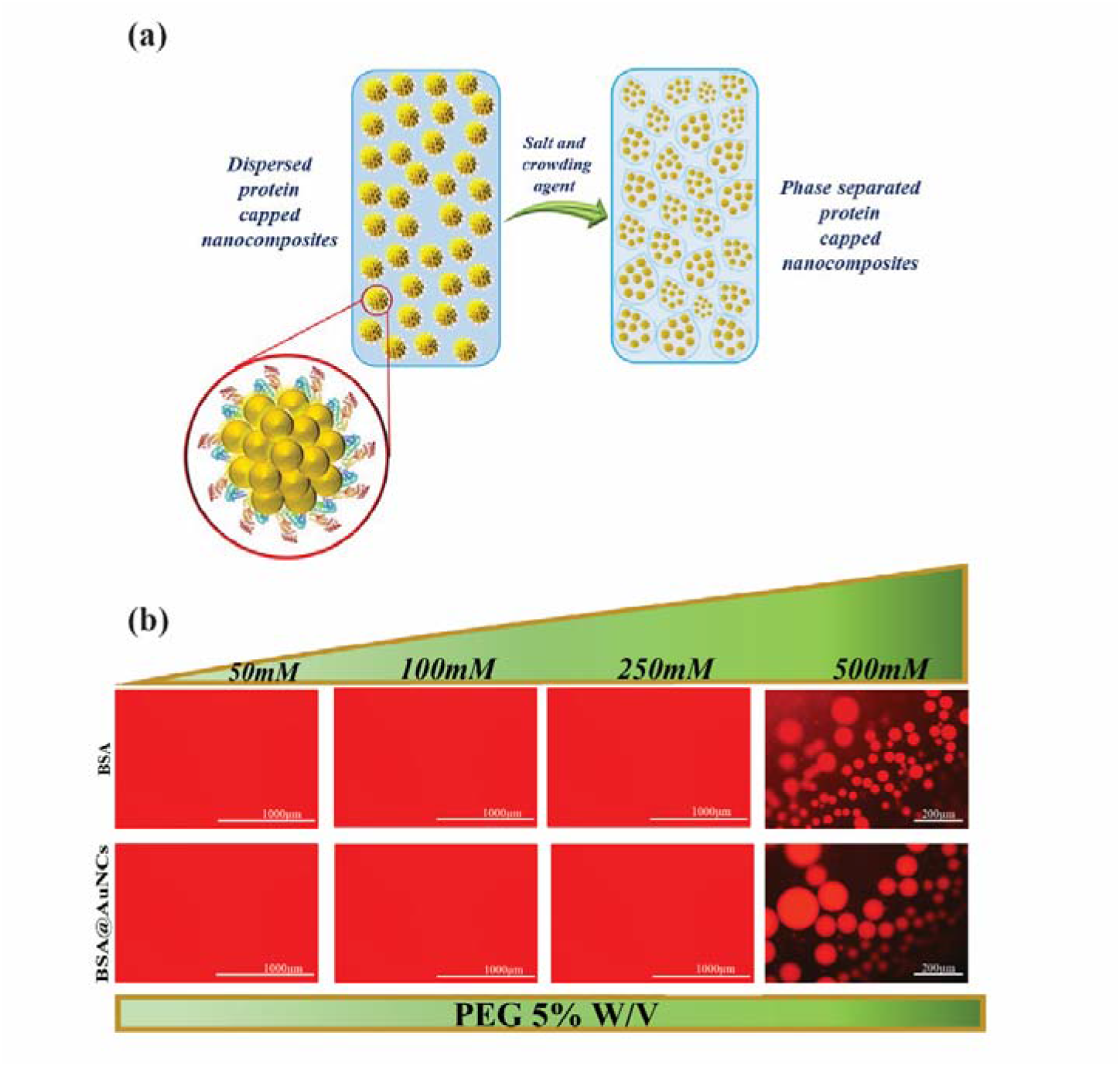
Phase separation of BSA-Au nanocomposites (a), Schematic illustration of phase separation of BSA-Au nanocomposites to liquid droplets in response to addition of salt and crowding agent, (**b, upper panel**) Microscopic visualization of the dynamics of BSA and BSA@AuNCs (**b, lower panel**) in the presence of 50-500 mM of sodium sulphate and crowding agent PEG-6000 (5%w/v). Scale bars represent 1000 μm for 50-250mM of sodium sulphate and 200 μm (500mM).

Next, we scanned different ionic strength with the kosmotropic salt sodium sulphate in the presence of polyethylene glycol (PEG-6000, 5% weight by volume) to induce LLPS in BSA, BSA@AuNCs and BSA@AuNPs. Both PEG and kosmotropes have been reported in the literature to aid the formation of LLPS ^[16]^. Texas-red labelled BSA is used in this experiment to assist the microscopic visualization of the event of LLPS. We observed that 500mM Na_2_SO_4_ in the presence of 5% w/v PEG6000 triggered the phase separation of BSA, BSA@AuNCs and BSA@AuNPs **(Fig 1b, Fig. S3)**. The fluid nature of the droplets is confirmed from the fusion events ( **Movie SV1** and **Movie SV2)**.

### Co-localisation of protein and gold metal inside liquid droplets

As we have used Texas-red labelled BSA, the obvious question next was if the observed droplets recruited BSA alone or the BSA coated nanocomposites. To clarify this, we designed a scattering-based light-microscopic assay. The underlying hypothesis for this experiment was based on the unique light scattering characteristic of gold nanoclusters ^[17]^. Differences in the intensity of scattered light collected from the liquid droplets of BSA and BSA@AuNCs would enable us to track the presence of gold nanoclusters inside droplets. Accordingly, we phase separated BSA@AuNCs and BSA individually into liquid droplets and performed imaging in bright-field microscopy. We used a 535 nm light source for scattering as the AuNCs have minimal absorbance at this wavelength. For imaging, we then drop-casted liquid droplets onto the glass coverslip and collected the elastically scattered light using bandpass filters (See scattering experiment detailed in materials and method). We noticed liquid droplets for both BSA@AuNCs and BSA proteins in the brightfield images, however, intense scattering was collected from the droplets containing BSA@AuNCs **(Fig. 2a,2b)** and not from the droplets made up of BSA alone, thus confirming the recruitment of the AuNCs inside the liquid droplets. Further, enhanced emission intensity observed in case of phase separated nanoclusters indicates their spatiotemporal confinement inside the liquid droplets ^[18,19]^ **(Fig. S4)**.

**Figure 2:**
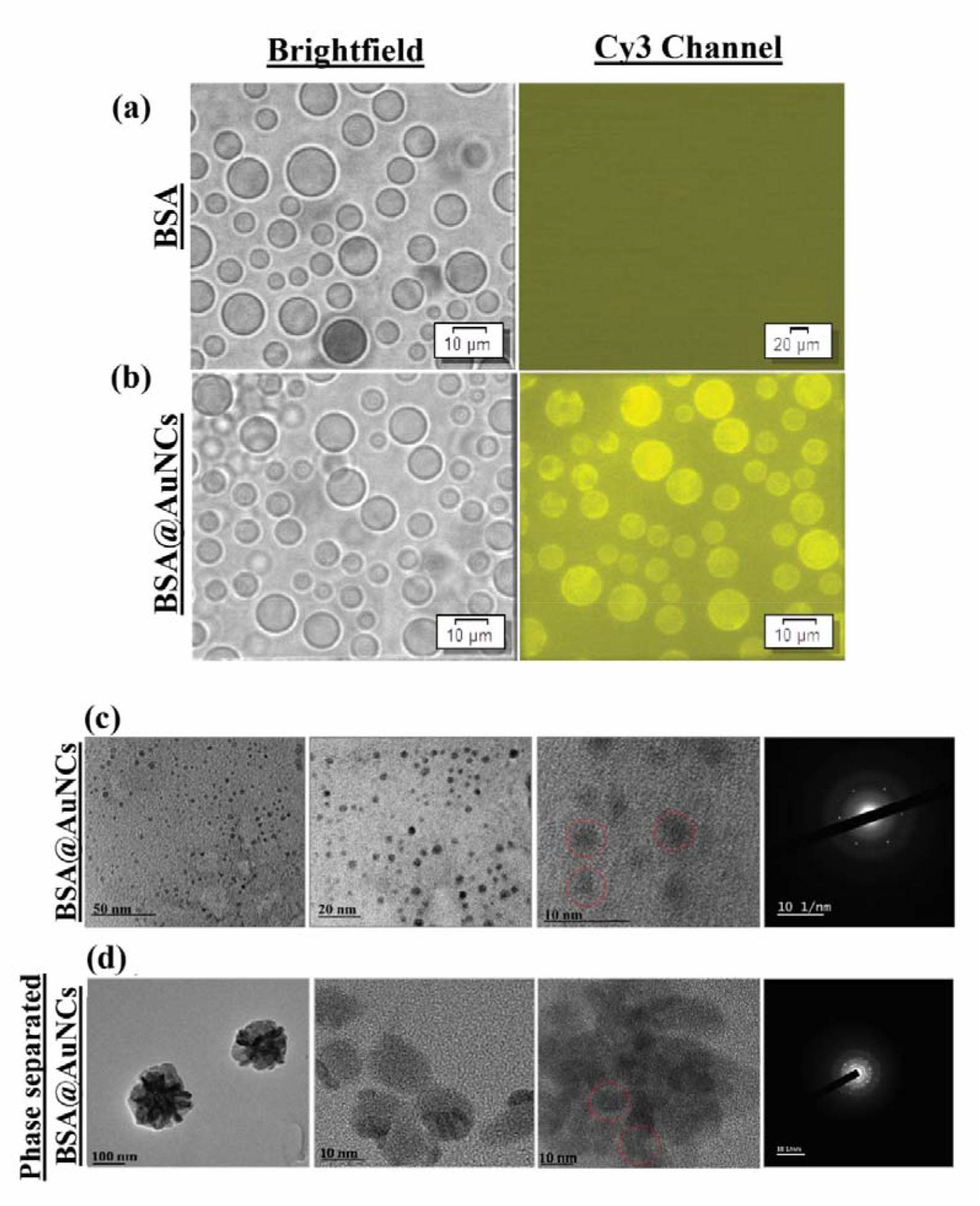
Structural elucidation of liquid droplets of BSA@AuNCs. Brightfield and scattering intensity images collected in Cy3 channel of liquid droplets formed by (a) BSA and (b) BSA@AuNCs. TEM images of (c) dispersed BSA@AuNCs and (d) condensates of BSA@AuNCs in the liquid droplets.

In the next set of experiments, we acquired transmission electron micrographs of BSA@AuNCs before and after LLPS **(Fig. 2c).** Post formation of the liquid droplets of BSA@AuNCs appeared as micron size assemblies of NCs compared to the discrete nature before the LLPS **(Fig. 2d**). The SAED patterns of BSA@AuNCs showed circular rings indicating dispersed protein capped NCs in solution **(Fig. 2c)** while phase-separated droplets feature diffusive rings. The diffusive rings indicate that the protein-nanoclusters in liquid droplets self-assemble to form bigger clusters **(Fig. 2d)**. Interestingly, the hydrodynamic diameter of the BSA@AuNCs in liquid droplets showed no significant difference from the dispersed condition. Thus, the AuNCs in liquid droplets are confined but not aggregated. The bigger aggregates in the electron micrographs are, however, an artifact of drying during the sample preparation.

### Enhanced catalysis in phase separated droplets of BSA-Au nanocomposites

Au nanocomposites serve as superior heterogeneous catalysis in several paradigms. We performed reduction and oxidation reactions using BSA@AuNCs and BSA@AuNPs as catalysts in the dispersed condition and phase separated environments.

The conversion of *p-nitrophenol* to *p*-aminophenol using sodium borohydride (NaBH_4_) as a reducing agent ^[20]^ is performed in the presence of BSA@AuNCs and LLPS BSA@AuNCs. The reaction is monitored by the gradual drop in the absorbance of *p-nitrophenol* at 400 nm. We noticed that the BSA@AuNCs in liquid droplets [*k*_*rate*_ = (**25.0 ± 0.4)** · **10^-3^ s^-1^**] drove the reaction one order faster than the dispersed BSA@AuNCs in solution [*k*_*rate*_ = (**3.0 ± 0.2)** · **10^-3^ s^-1^**] (**Fig. 3a).** The near ten-fold enhancement in the reaction-rate in the phase separated environment is most-likely due to the increased local concentration of substrate and catalysts within the liquid droplets. BSA@AuNPs also showed seven-fold faster rate of reaction in condensed form [*k*_*rate*_ = (**20±0.3)**· **10^-3^ s^-1^**] than the nanoparticles in dispersed phase [*k*_*rate*_ = (**2.8±0.6)** · **10^-3^s^-1^**] **(Fig. S5)**. Solute-solute and solvent-solvent interactions facilitate the phase separation in solution. Thus, it is expected that the condensed phase of BSA@AuNCs in solution will remain stable for a longer time. We observed that BSA@AuNCs condensed phases remain stable in solution **(Fig. S6)** and show optimum catalytic activity upto five days of measurements **(Fig. 3b).** Overall, our experimental results demonstrate that the catalytic droplets not only enhance the catalytic efficiency of the recruited nanoclusters by proximal interactions and efficient substrate channeling, but the like solvent-solvent and solute-solute interactions retain the catalysts active in solution for a significantly longer time.

**Figure 3:**
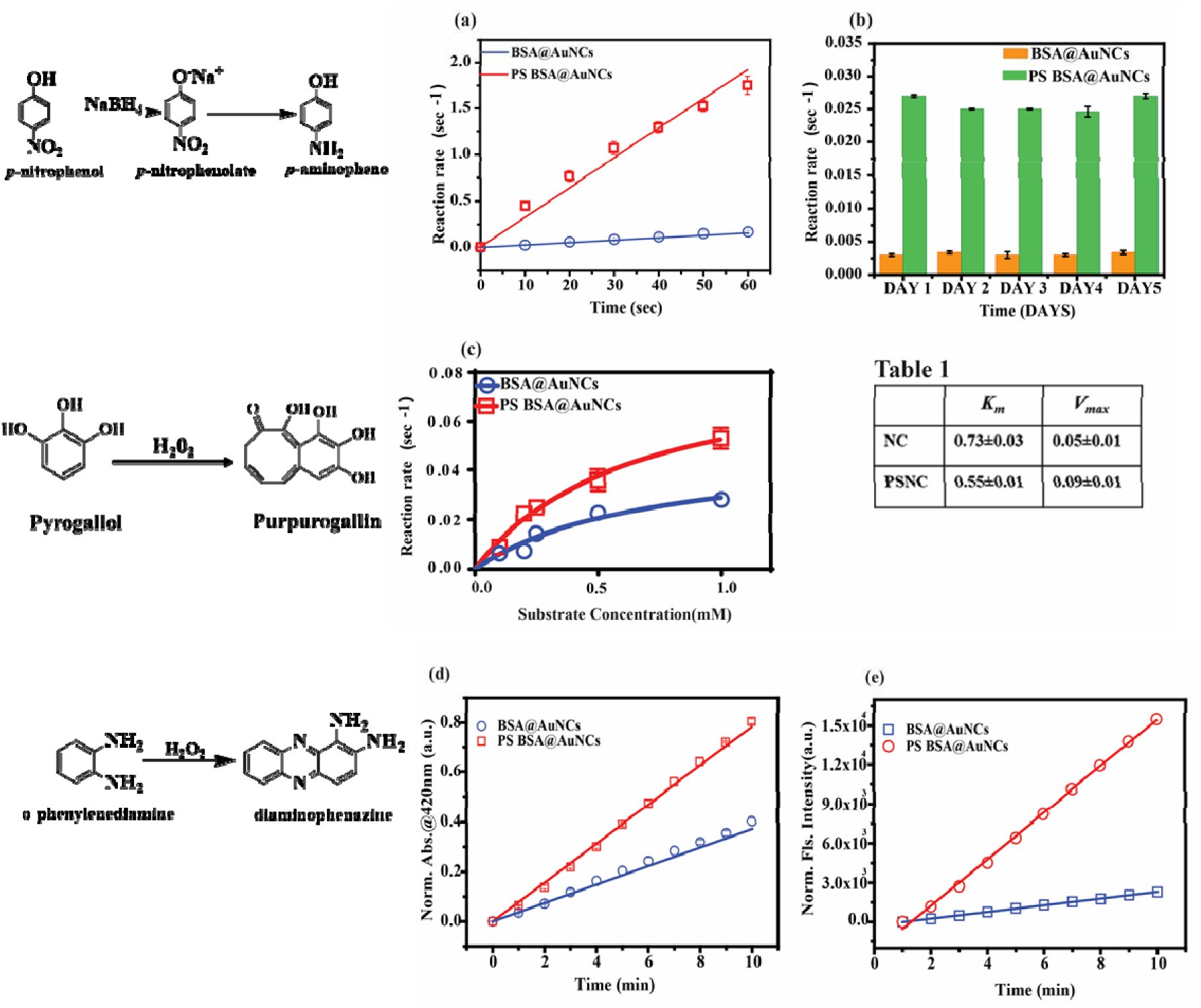
Catalytic efficiency of liquid droplets of BSA @AuNCs. (a) Rate of reaction for reduction of p-nitrophenol using BSA@AuNCs in dispersed (blue line) and in phase separated condensate form (PS BSA@AuNCs, red line), (b) Dispersion stability of BSA@AuNCs in liquid condensate form for five consecutive days where the activity is monitored from the reduction of p-nitrophenol. (c) Michaelis –Menten kinetics fit for oxidation reaction of pyrogallol to purpurogallin using BSA@AuNCs in dispersed (blue line) and phase separated BSA@AuNCs (PS BSA@AuNCs, red line). Table 1 indicates the K_m_ and V_max_ values for BSA@AuNCs in dispersed solution (NC) and in phase separated form (PSNC). (d) Rate of reaction of DAP formation using BSA@AuNCs in dispersed (blue line) and in phase separated form (red line), (e) Rate of increase in fluorescence intensity at 565nm with time using BSA@AuNCs in dispersed solution (blue line) and in phase separated form (red line).

We next studied the peroxidase mimicking properties of the BSA@AuNCs^[21]^ by testing the oxidation of substrate pyrogallol to purpurogallin. We computed the maximal velocity (*V*_*max*_) and Michaelis-Menten constant (*K*_*m*_) in dispersed and phase separated conditions using Michaelis–Menten equation (**Fig. 3c**). A lower *K*_*m*_ and similar *V*_*max*_ is obtained in the case of phase separated BSA@AuNCs in comparison to the dispersed BSA@AuNCs. A lower *K*_*m*_ is indicative of higher substrate affinity with the catalyst while the *V*_*max*_ indicates maximum rate of the enzyme catalysis **(Table 1)**. The lower *K*_*m*_ of reaction in the phase separated form may be due to enhanced proximity of substrate and catalyst within the liquid condensates formed in the phase separation event ^[12]^.

In another assay, we also compared the rate of oxidation reaction of o-phenylenediamine (OPD) to a fluorescent product diaminophenazine (DAP). Kinetics of reaction in bulk and phase separated form is shown in supplementary **(Fig.S7).** DAP absorbs at 420 nm and emits at 565 nm. We observed a faster rate of oxidation in phase separated nanoclusters, [*k*_*rate*_ = (**8± 0.086)** · **10^-2^ min^-1^**] than in dispersed state [*k*_*rate*_ = (**4.0± 0.025)** · **10^-2^ min^-1^**] **[Fig.3d]**. Further, enhanced steady-state fluorescence intensity in phase separated nanoclusters suggest more product formation in phase separated environments **(Fig. 3e)**.

The three discrete chemical catalysis reactions described above show that the phase separated nanoclusters lead to an increased rate of reaction by concentrating substrate and catalyst within the liquid droplets. Macromolecular crowding results in an increase in the effective concentration of reactants and catalysts and their association rates within the droplets ^[22]^. Our results are at par with the results reported for the higher catalytic activity of enzymes upon condensing in a liquid phase from a solution phase ^[23]^. Such an enhanced catalytic activity is, however, reported for the first time for phase separated inorganic nanoconjugates.

### Phase separation and catalytic reactions of α-synuclein coated gold nanoparticles

Does the phase separation of protein-based nanocomposites depend on the native structure of the protein? To monitor, we synthesized Au nanoparticles coated with an intrinsically disordered protein, α-synuclein. Absorbance spectra recorded for α-synuclein coated Au nanoparticles (α-synAuNPs) showed peak at 520 nm correspond to the surface plasmon resonance peak of Au nanoparticles and a shoulder peak at 275 nm for the tyrosine residues present in α-synuclein **(Fig.S8).** DLS data showed the increase in the hydrodynamic diameter **(Fig.S9a)** and decrease in the zeta potential in case of α-synAuNPs **(Fig.S9b)** that confirms the interaction of positively charged N-terminal of α-synuclein with negatively charged citrate capped Au nanoparticles. Further, we observed that changing the ionic strength of solution in presence of a crowding agent leads to the phase separation of α-synAuNPs. 500mM of Na_2_SO_4_ and 5% w/v of PEG-6000 forms the liquid droplets of α-synAuNPs **(Fig. S10)**. We synthesize the nanoparticles with Alexa-488 labeled α-synuclein and thus, the green coloured droplets.

We subsequently determine the change in the catalytic properties of α-synAuNPs before and after inducting the phase separation. Rates of *p*-nitrophenol reduction in dispersed state and in liquid droplets are [*k*_*rate*_ = (**7± 0.3)** · **10^-3^ s^-1^**] and [*k*_*rate*_ = (**61 ± 0.1)** · **10^-3^ s^-1^**], respectively **(Fig. 4a)**. The phase separation of α-synAuNPs fastened the reduction reaction by nearly 9 folds. On performing the peroxidase mimicking enzyme activity of α-synAuNPs, we inferred the lower K_*m*_ and comparable V_*max*_ in the case of phase separated α-synAuNPs **(Fig. 4b, Table 2)**, once again indicating a higher catalytic proficiency of protein-capped nanocatalysts in a phase separated environment in comparison to the dispersed state.

**Figure 4:**
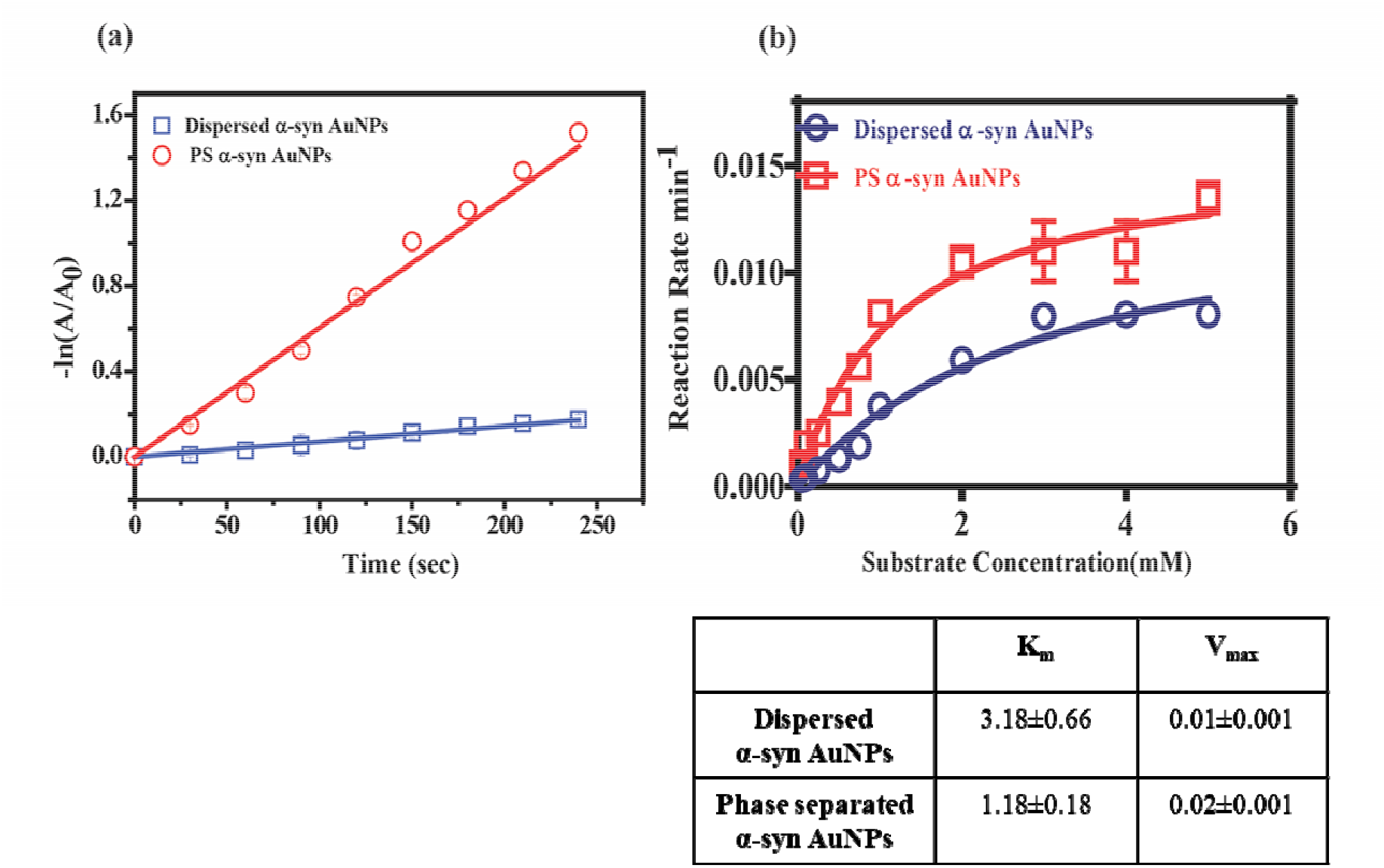
Catalytic efficiency of liquid droplets of α*-syn@AuNPs-*. (a) Rate of reaction for reduction of *p*-nitrophenol using α-syn AuNPs in dispersed (blue line) and in phase separated condensate form (PS α-syn AuNPs, red line), (b) Michaelis –Menten kinetics fit for oxidation reaction of pyrogallol to purpurogallin using α-syn AuNPs in dispersed (blue line) and phase separated α-syn AuNPs (PS α-syn AuNPs, red line), Table 2 indicates the K_m_ and V_max_ values for α-syn AuNPs in dispersed solution and in phase separated form.

### Phase separation occurs irrespective of the size and type of protein-nanocatalysts

Our next objective was to explore if protein-metal nanocomposites of any sizes can undergo LLPS. So, we checked the phase separation of BSA-CuPO_4_ nanoflowers that are micron in size **(Fig. S11)**. We induced phase separation for this nanocomplex, as before. As expected, BSA-CuPO_4_ nanoflowers undergo phase separation and form liquid droplets. Peroxidase activity of nanoflowers in homogenous solution and in phase separation was compared by calculating *K*_*m*_ and *V*_*max*_ of the kinetic reaction using Michaelis-Menten equation **(equation 2)**. Nanoflowers in phase separated form show lower *K*_*m*_ and comparable *V*_*max*_ **(Fig.S12)** in comparison to homogenous solution of nanoflowers (**Table S1)**. Lower *K*_*m*_ values indicate the high affinity of substrate and catalysts within the condensate phase.

## Discussion

Confining inorganic catalysts within well-defined structures of MOFs, COFs for enhanced catalytic activities are established in heterogeneous catalysis ^[2]^. However, confining the nanocomposites in membrane-less droplets with free-diffusivity is still less explored. Phase separation has emerged as a potential mechanism to accelerate the biochemical reactions by providing the spatiotemporal confinement of catalysts ^[24]^. A considerable number of studies have explained the importance of phase separation in increasing enzymatic reaction rate ^[25]^. Several reports have illustrated that biological proteins quickly respond to changes in their microenvironment and undergo LLPS to form membrane-less organelles. These studies motivated us to study the effect of microenvironment on the phase separation behavior of protein-based nanomaterials. In our work, we demonstrate to prepare a stimulus responsive protein-metal nanohybrid system that can undergo liquid-liquid phase separation. We used gold nanostructure as a nanocatalyst that exhibits excellent catalytic properties. We capped the nanostructures with proteins and increased its catalytic proficiency by inducing phase separation. The phase separation of the protein-Au NC occurs irrespective of the native structure of the protein or the size of the synthesized nanocomposites. This strengthens the idea that the process of phase separation observed in our study is environment dependent and is not influenced by the nature or size of the nanomaterial under investigation.

## Materials

All the chemicals unless mentioned were procured from Sigma-Aldrich. Copper chloride dihydrate and sodium sulphate were purchased from HiMedia. Ortho-phenylenediamine (OPD) was procured from TCI chemicals. Construct of α-synuclein was kindly provided by Prof. Samrat Mukhopadhyay, IISER Mohali. All other reagents were of analytical grade and used as received. Ultrapure water was used throughout the experiments.

## Methodology

### Synthesis and characterisation of BSA-Au nanocomposites

We synthesized BSA capped gold nanoclusters (BSA@AuNCs) and BSA capped gold nanoparticles (BSA@AuNPs) according to the method reported by Xie ^[26]^ and Matei *et al* ^[27]^ with some modifications. For BSA@AuNCs, 1 ml of 10 mg/ml of dialysed BSA protein solution was mixed with HAuCl_4_ solution (10mM). Reaction mixture was stirred vigorously at 37°C for half an hour followed by addition of 200μl of 1M NaOH. Reaction was kept at stirring for next 8-10 hours till the development of dark brown colored solution. Completion of reaction was confirmed by recording the absorption and fluorescence spectra of BSA@AuNCs using Cary UV-Vis compact peltier spectrophotometer (Agilent, USA) and Spectrofluorimeter FS5 (Edinburgh Instruments, UK) respectively. Prepared Nanoclusters were dialysed using 3.5 KDa dialysis membrane against distilled water for continuous 48 hours to make the pH of nanoclusters neutral and stored at 4°C for further use. Further, size of nanoclusters was confirmed using a ZetaSizer Nano ZSP (Malvern Instruments, UK). The scattered intensity was measured at a backscattered angle of 173°. For each sample, three readings were recorded. To understand the morphology of the BSA@AuNCs, TEM images were obtained by drop casting 10 μl of 0.5 mg/ml nanocluster on glow discharged carbon coated TEM grids. The grids were then washed with Milli-Q water and dried using desiccator for 24 hours at room temperature. TEM imaging was done by using JEM 2100 TEM (JEOL, USA) operated at 200 kV.

To prepare BSA@AuNPs, we mixed BSA (5mg/ml) with HAuCl_4_ solution (100μM). Solution was vigorously stirred at room temperature. In order to optimize the NaBH_4_ concentration, various amounts of NaBH_4_ were added to the solution. The pink-red solution of BSA@AuNPs were prepared using 1mM concentration of NaBH_4._ Further, we recorded its UV-Vis absorption spectra using Cary UV-Vis compact peltier spectrophotometer (Agilent, USA)

### Phase separation of BSA-Au nanocomposites

To visualize the effect of changing ionic strength on BSA and BSA-Au nanocomposites, we performed fluorescence microscopy experiments. We labelled the BSA with TEXAS Red dye and removed the unbound dye by extensive dialysis against water for continuous 48 hours. We prepared samples having 1mg/ml of labelled BSA incubated with increasing sodium sulphate concentration in the range from 50mM-500mM followed by addition of 5% w/v crowding agent, PEG-6000. Next, we performed the fluorescence microscopic study of BSA@AuNCs and BSA@AuNPs prepared using texas-red labelled BSA. To prepare samples we used 1mg/ml of nanocomposite incubated with sodium sulphate (50-500mM) and 5% w/v of PEG-6000. Prepared samples were dropcasted on the microscope slides and visualized using a 10X objective lens in the red channel of Cytation-5 Cell imaging Multi-Mode Reader (BioTek Instruments).

### Co-localisation of protein and gold metal inside liquid droplets

To check the co-localisation of protein and gold metal inside liquid droplets, we designed a scattering-based light-microscopic assay. The design of the experiment was based on the hypothesis that the presence of metal ions in the droplets would scatter more light in comparison to a protein liquid droplet. For this experiment, samples were prepared by incubating 1mg/ml of BSA and BSA@AuNCs with 500mM Na_2_SO_4_ AND 5%w/v of PEG-6000 differently. The samples having phase separated liquid droplets were drop casted onto the glass coverslip and placed under 40X objective lens of inverted fluorescence microscope (IX83 P2ZF Inverted microscope equipped with IX83 MITICO TIRF illuminator). The droplets were excited using Cy3 channel (excitation wavelength 535 nm) with a laser intensity of 25% at an exposure time of 440 ns. Scattering was collected at Cy3 channel as metal nanoparticles do not emit at this wavelength. To check the change in morphology after phase separation of BSA@AuNCs as liquid condensates, we performed the TEM imaging of phase separated BSA@AuNCs. TEM images were obtained by drop casting 10 μl of sample having 1mg/ml of BSA@AuNCs mixed with 500mM sodium sulphate and 5% w/v PEG-6000 on to carbon coated TEM grids. The grids were then washed with Milli-Q water and dried using a desiccator for 24 hours at room temperature. TEM imaging was done by using JEM 2100 TEM (JEOL, USA) operated at 200 kV.

### Synthesis of BSA-Copper phosphate nanoflowers

BSA-Copper phosphate nanoflowers were prepared according to the latest publication from our lab by Kaur *et al* ^[28]^, incubating BSA solution (5μM) with CuCl_2_ dihydrate (500μM) in phosphate buffer saline pH 7.4. Sample was kept on stirring at room temperature for continuous 24 hours followed by centrifugation of the sample at 10,000rpm for ten minutes. Pellet was extensively washed with water to remove the excess salts. Further, we performed the Scanning electron microscopy to confirm the formation and morphology of synthesized BSA-Copper phosphate nanoflowers.

### Catalytic activity of BSA-metal nanocomposites

We checked the catalytic potential of prepared BSA@AuNCs and BSA@AuNPs by performing oxidation and reduction of aromatic compounds in dispersed and phase separated form. Reduction of *p-nitrophenol* to *p-aminophenol* was performed using BSA@AuNCs and BSA@AuNPs in dispersed and phase separated form. The concentration of both the systems was normalized by keeping the concentration of BSA constant. 1mg/ml of BSA@AuNCs and BSA@AuNPs was mixed with 100μM of *p-nitrophenol*. Initial absorbance spectra of solution were recorded using TECAN INFINITE MPLEX reader followed by addition of 100mM of reducing agent sodium borohydride. Decrease in absorbance at 400 nm corresponding to reduction of *p-nitrophenol* was monitored for next ten minutes in kinetics mode at each ten second time interval. Similar catalytic reduction was performed using BSA@AuNCs in phase separated form, having 1mg/ml of nanocluster, 500mM sodium sulfate and 5%w/v PEG-6000 mixed with 100μM of *p-nitrophenol*. Further, we calculated the rate of reaction for BSA@AuNCs in dispersed and phase separated form, from the slope of the curve by fitting the data to first order decay kinetics equation (**equation 1**) using OriginPro 8.6 software.

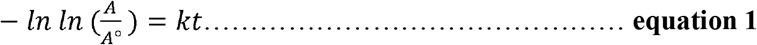

(A-Absorbance value at certain time points, A_°_-Initial absorbance value, k-rate of reaction, t-time in seconds).

### Peroxidase activity

To check the peroxidase activity of BSA@AuNCs and BSA-CuPO_4_ nanoflowers, oxidation of pyrogallol to purpurogallin was done in dispersed and in phase separated form. Reaction mixture was having 1mg/ml of BSA@AuNCs and 20μl of nanoflowers of concentration 5μM separately and mixed with different concentrations of pyrogallol (100μM-1000μM) followed by addition of 0.1M H_2_O_2._ Production of purpurogallin was tracked by observing increase in absorbance at 420 nm using TECAN INFINITE MPLEX plate reader at kinetics mode. Reading was recorded at a time interval of one minute for a complete one hour. Next, the experiment was done using both the nanocomposites in a phase separated state with the same concentrations of pyrogallol as explained above with 500mM sodium sulfate and 5%w/v of PEG-6000 followed by addition of 0.1M H_2_O_2._ Further, rate of reaction at different pyrogallol substrate concentration was plotted and *K*_*m*_ and *V*_*max*_ of the reaction was calculated after fitting the data in Michaelis-Menten kinetics equation (**equation 2)** given below using GraphPad Prism 6 software for BSA@AuNCs in both the phases.

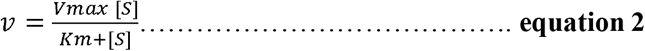

*v-* velocity of reaction, V_max_*-* maximum rate of reaction achieved by the system, [S]-Substrate concentration, K_m_-Michaelis-Menten constant.

Peroxidase activity of prepared BSA@AuNCs was also checked against oxidation of another peroxidase substrate *o*-phenylenediamine to yellow colored product diaminophenazine. Peroxidase assay was performed using 1mg/ml of nanoclusters solution with 0.2mM OPD and 1mM H_2_0_2_. Formation of product DAP was tracked by recording absorbance and PL spectra of the reaction mixture in a steady-state kinetic assay of 30 minutes with 1 minute time interval. For PL spectra sample was excited at 400 nm and an increase in emission intensity of the product at 565 nm was tracked for continuous 30 minutes. Further, we performed the peroxidase kinetic assay after inducing liquid-liquid phase separation of the BSA@AuNCs using 500mM sodium sulphate and 5%w/v PEG-6000. We fitted the absorbance and fluorescence spectral data to calculate the rate of reaction in both conditions using the first order kinetic equation represented as **equation 1** above.

### Synthesis and phase separation of α-synuclein coated gold nanoparticles

We purified the α-synuclein protein according to the protocol by Jain *et al* ^[29]^. Overnight grown primary culture was transferred to the fresh media followed by addition of 0.8mM IPTG and incubated at 37°C for 4 hours. Cells were harvested by centrifuging the culture at 8000 rpm for ten minutes. Pellet was resuspended in the lysis buffer and boiled at 90°C for 30 minutes. Lysed cells were centrifuged at 11,000rpm followed by addition of 10% streptomycin sulfate and glacial acetic acid to the supernatant. Further, a saturated solution of ammonium sulphate was used to precipitate out the protein. To synthesize α-syn AuNPs, we first synthesize the citrate capped gold nanoparticles using trisodium citrate as a reducing agent ^[30]^. Gold chloride solution of 500μM concentration was heated to boil followed by addition of 10μl of 5%w/v of trisodium citrate. Solution was continuously stirred for next one hour till the color of the solution was changed to wine-red. Further, we incubated the citrate capped gold nanoparticles with 250⎧M of purified α-synuclein protein for 24 hours with continuous stirring at 4°C. Sample was centrifuged and repetitively washed with water. Further, we resuspend the pellet in 500⎧l of distilled water to perform the further experiments. Conjugated protein concentration was calculated from the absorbance spectra of α-syn AuNPs. To phase separate α-syn AuNPs we incubated the α-syn AuNPs with 500mM sodium sulphate and 5% w/v of PEG-6000. We dropcasted the sample on a microscopic slide and observed the formation of liquid droplets using upright fluorescence microscope OLYMPUS BX53 (OLYMPUS; USA).

### Catalytic activity of α-synuclein coated gold nanoparticles

We performed the oxidation and reduction reaction of aromatic compounds using α-synuclein coated gold nanoparticles. Reduction of *p*-nitrophenol to *p*-aminophenol is performed using 70μM of protein coated gold nanoparticles with 50⎧M of PNP followed by addition of 4mM of NaBH_4_. Decrease in absorbance at 400 nm was noted in a kinetic cycle of 15 minutes with 30 second time interval. Oxidation of pyrogallol to purpurogallin was also performed using the same concentration of α-synuclein coated gold nanoparticles. Reaction was performed at different concentrations of pyrogallol ranging from 100mM-3mM in dispersed and phase separated states. Formation of purpurogallin was observed by recording an increase in absorbance corresponding to 420 nm. We calculated the K_*m*_ and V_*max*_ of the reaction after fitting the data in Michaelis-Menten equation using GraphPad Prism software.

## Conclusion

Confinement presumes to enhance the catalytic activity^[31,32]^. Unfortunately, the physical barriers in the confinement often separate the catalysts from solutions and introduce heterogeneity in the system. Conversely, homogeneous catalysis has experimentally demonstrated far superiority in activity than the equivalent heterogeneous systems. Thus, with an overarching objective to obtain a homogeneous confinement of catalysts in solution, we induced phase separations of nano-catalysts into liquid droplets within solution and performed their catalytic activity. Here, we demonstrated a strategy to condense a protein-nanocomposite in a liquid phase from a solution phase by altering the ionic strength of the solution along with the use of crowding agents. We observed that confinement of catalysts is responsive to change in the microenvironment of solution and it is independent of the native structure of the protein wrapped outside the nanomaterial. We next compared the catalytic activities of these bio-nanohybrids dispersed in solution and condensate phase. Liquid condensates of nanocomposite showed nearly one order of magnitude higher catalytic activity than the dispersed phase. This demonstration may be extended to other hybrid catalytic systems for the development of novel stimuli responsive novel bio-inorganic systems for enhanced catalytic properties^[33]^. Overall, our findings pave the way to use the protein based nanohybrids in a phase separated form like other biological enzymes to enhance their catalytic potential.

---

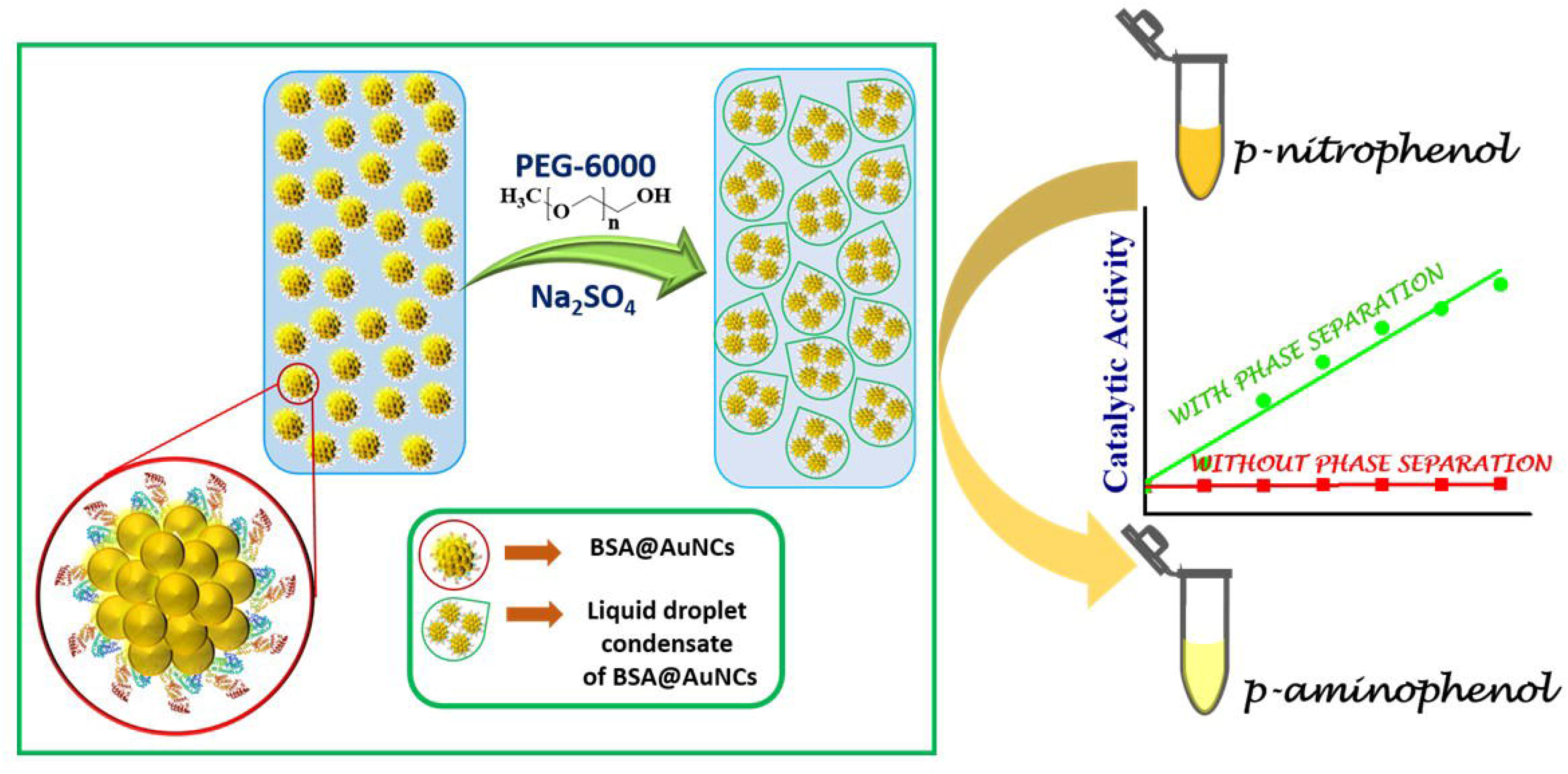

## References

1. Cao S, Tao FF, Tang Y, Li Y, Yu J. Size-and shape-dependent catalytic performances of oxidation and reduction reactions on nanocatalysts. Chemical Society Reviews. 2016;45(17):4747–65.

2. Gao G, Xi Q, Zhang Y, Jin M, Zhao Y, Wu C, Zhou H, Guo P, Xu J. Atomic-scale engineering of MOF array confined Au nanoclusters for enhanced heterogeneous catalysis. Nanoscale. 2019;11(3):1169–76.

3. Li H, Xiao J, Fu Q, Bao X. Confined catalysis under two-dimensional materials. Proceedings of the National Academy of Sciences. 2017 Jun 6;114(23):5930–4.

4. Dhakshinamoorthy A, Asiri AM, Garcia H. Catalysis in confined spaces of metal organic frameworks. ChemCatChem. 2020 Oct 6;12(19):4732–53.

5. Dhakshinamoorthy A, Li Z, Garcia H. Catalysis and photocatalysis by metal organic frameworks. Chemical Society Reviews. 2018;47(22):8134–72.

6. Gao C, Lyu F, Yin Y. Encapsulated metal nanoparticles for catalysis. Chemical Reviews. 2020 Jun 25;121(2):834–81.

7. Hyman AA, Weber CA, Jülicher F. Liquid-liquid phase separation in biology. Annual review of cell and developmental biology. 2014 Oct 6;30:39–58.

8. Mao YS, Zhang B, Spector DL. Biogenesis and function of nuclear bodies. Trends in Genetics. 2011 Aug 1;27(8):295–306.

9. Cioce M, Lamond AI. Cajal bodies: a long history of discovery. Annu. Rev. Cell Dev. Biol.. 2005 Nov 10;21:105–31.

10. Buchan JR. mRNP granules: assembly, function, and connections with disease. RNA biology. 2014 Aug 3;11(8):1019–30.

11. Dzuricky M, Rogers BA, Shahid A, Cremer PS, Chilkoti A. De novo engineering of intracellular condensates using artificial disordered proteins. Nature chemistry. 2020 Sep;12(9):814–25.

12. Peeples W, Rosen MK. Mechanistic dissection of increased enzymatic rate in a phase-separated compartment. Nature chemical biology. 2021 Jun;17(6):693–702.

13. Ray S, Singh N, Kumar R, Patel K, Pandey S, Datta D, Mahato J, Panigrahi R, Navalkar A, Mehra S, Gadhe L. α-Synuclein aggregation nucleates through liquid–liquid phase separation. Nature chemistry. 2020 Aug;12(8):705–16.

14. Van den Berg B, Wain R, Dobson CM, Ellis RJ. Macromolecular crowding perturbs protein refolding kinetics: implications for folding inside the cell. The EMBO journal. 2000 Aug 1;19(15):3870–5.

15. Qian H, Zhu M, Wu Z, Jin R. Quantum sized gold nanoclusters with atomic precision. Accounts of chemical research. 2012 Sep 18;45(9):1470–9.

16. Mohammadi P, Jonkergouw C, Beaune G, Engelhardt P, Kamada A, Timonen JV, Knowles TP, Penttila M, Linder MB. Controllable coacervation of recombinantly produced spider silk protein using kosmotropic salts. Journal of colloid and interface science. 2020 Feb 15;560:149–60.

17. Priest L, Peters JS, Kukura P. Scattering-based Light Microscopy: From Metal Nanoparticles to Single Proteins. Chemical Reviews. 2021 Sep 29;121(19):11937–70.

18. Huang Z, Wang M, Guo Z, Wang H, Dong H, Yang W. Aggregation-enhanced emission of gold nanoclusters induced by serum albumin and its application to protein detection and fabrication of molecular logic gates. ACS omega. 2018 Oct 8;3(10):12763–9.

19. Xia M, Sui Y, Guo Y, Zhang Y. Aggregation-induced emission enhancement of gold nanoclusters in metal–organic frameworks for highly sensitive fluorescent detection of bilirubin. Analyst. 2021;146(3):904–10.

20. Lin C, Tao K, Hua D, Ma Z, Zhou S. Size effect of gold nanoparticles in catalytic reduction of p-nitrophenol with NaBH4. Molecules. 2013 Oct;18(10):12609–20.

21. Wang S, Liu P, Qin Y, Chen Z, Shen J. Rapid synthesis of protein conjugated gold nanoclusters and their application in tea polyphenol sensing. Sensors and Actuators B: Chemical. 2016 Feb 1;223:178–85.

22. Jiang L, Chen X, Lu N, Chi L. Spatially confined assembly of nanoparticles. Accounts of chemical research. 2014 Oct 21;47(10):3009–17.

23. Küchler A, Yoshimoto M, Luginbühl S, Mavelli F, Walde P. Enzymatic reactions in confined environments. Nature nanotechnology. 2016 May;11(5):409–20.

24. Kojima T, Takayama S. Membraneless compartmentalization facilitates enzymatic cascade reactions and reduces substrate inhibition. ACS applied materials & interfaces. 2018 Sep 4;10(38):32782–91.

25. Krainer G, Welsh TJ, Joseph JA, Espinosa JR, Wittmann S, de Csilléry E, Sridhar A, Toprakcioglu Z, Gudiškytė G, Czekalska MA, Arter WE. Reentrant liquid condensate phase of proteins is stabilized by hydrophobic and non-ionic interactions. Nature communications. 2021 Feb 17;12(1):1–4.

26. Xie J, Zheng Y, Ying JY. Protein-directed synthesis of highly fluorescent gold nanoclusters. Journal of the American Chemical Society. 2009 Jan 28;131(3):888–9.

27. Matei I, Buta CM, Turcu IM, Culita D, Munteanu C, Ionita G. Formation and stabilization of gold nanoparticles in bovine serum albumin solution. Molecules. 2019 Jan;24(18):3395.

28. Kaur H, Bari NK, Garg A, Sinha S. Protein morphology drives the structure and catalytic activity of bio-inorganic hybrids. International Journal of Biological Macromolecules. 2021 Apr 15;176:106–16.

29. Jain N, Bhasne K, Hemaswasthi M, Mukhopadhyay S. Structural and dynamical insights into the membrane-bound α-synuclein. PloS one. 2013 Dec 20;8(12):e83752.

30. Mohammadi S, Nikkhah M. TiO2 nanoparticles as potential promoting agents of fibrillation of α-synuclein, a parkinson’s disease-related protein. Iranian journal of biotechnology. 2017;15(2):87.

31. Grommet AB, Feller M, Klajn R. Chemical reactivity under nanoconfinement. Nature nanotechnology. 2020 Apr;15(4):256–71.

32. Li X, Wang N, Bao X, Li Q, Li J, Xie YB, Ji S, Yuan J, An QF. Nano-confinement-inspired metal organic framework/polymer composite separation membranes. Journal of Materials Chemistry A. 2020;8(33):17212–8.

33. Bari NK, Kumar G, Bhatt A, Hazra JP, Garg A, Ali ME, Sinha S. Nanoparticle fabrication on bacterial microcompartment surface for the development of hybrid enzyme-inorganic catalyst. ACS Catalysis. 2018 Jul 24;8(9):7742–8.

