## Supplementary information for "Barrier-free liquid condensates of nanocatalysts as effective concentrators of catalysis"

**Supplementary Figures**

(a) **(b)**

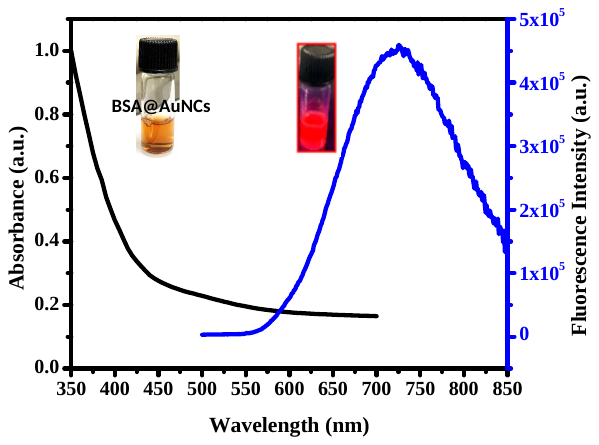

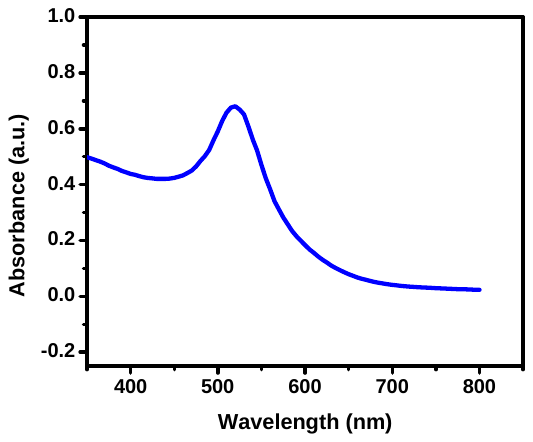

**(c)	(d)**

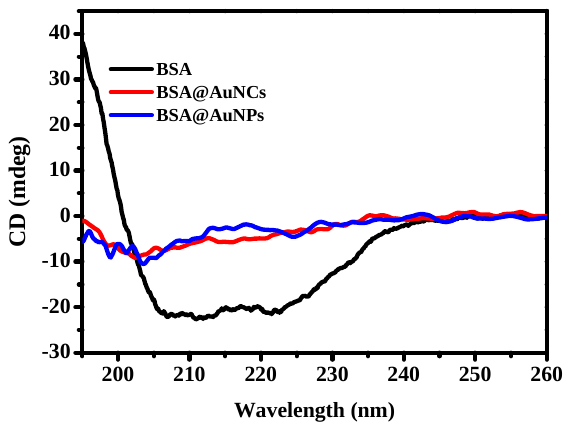

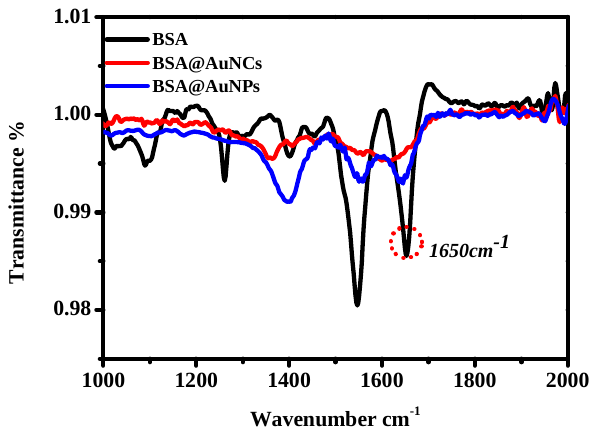

**Figure S1:** (a) Absorbance and fluorescence spectra (excitation at 450nm) of BSA@AuNCs. Inset shows the suspensions of BSA@AuNCs in stray light and under UV-lamp, (b) Absorbance spectra of BSA@AuNPs, (c) CD spectra of native structure of BSA, BSA@AuNCs and BSA@AuNPs showing decrease in α-helix content of BSA in BSA@AuNCs, (d) FTIR spectra of BSA, BSA@AuNCs and BSA@AuNPs showing disappearance of peaks correspond to α-helix of BSA in BSA@AuNCs.

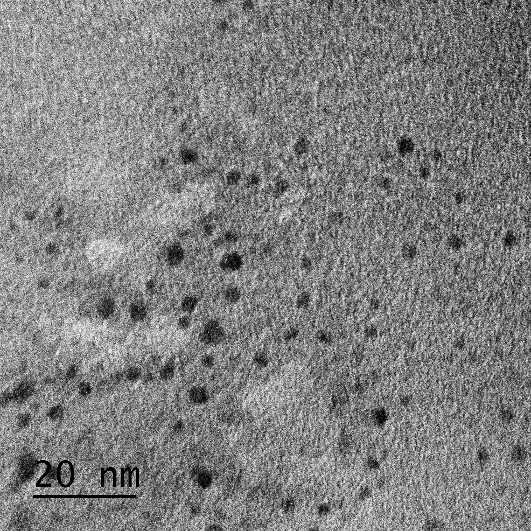

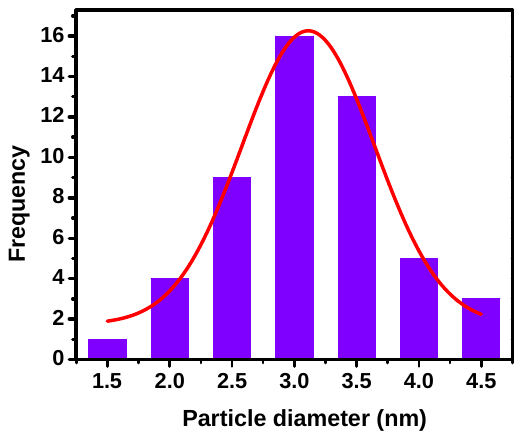

**Figure S2:** TEM image of BSA@AuNCs, showing dispersed nanoclusters of size 3.0±0.5nm, histogram shows the frequency distribution plot of average size of fifty particles.

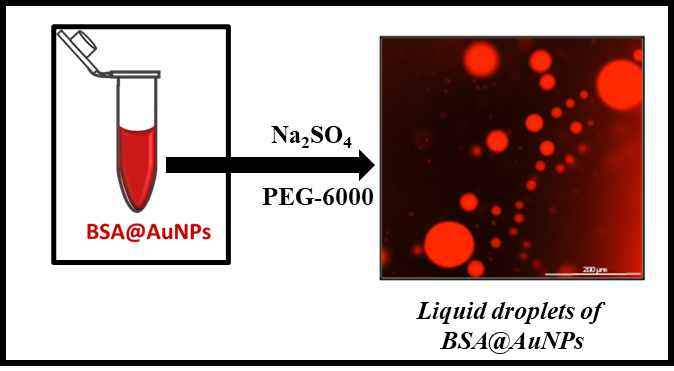

**Figure S3:** Absorbance spectra of BSA capped gold nanoparticles, shows sharp absorbance at 520nm, Inset shows the wine colored BSA@AuNPs. Addition of sodium sulphate and PEG-6000 forms liquid droplets of BSA@AuNPs

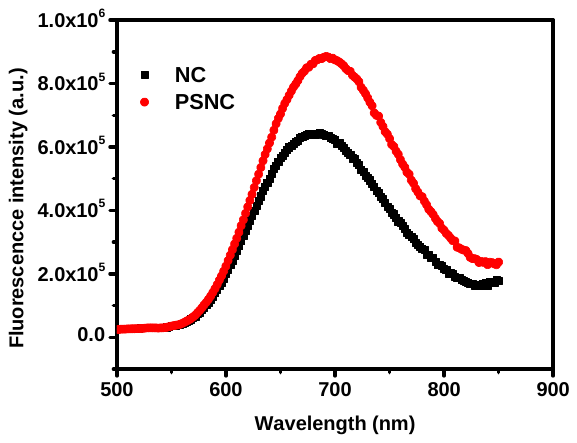

**Figure S4:** Fluorescence emission spectra of BSA@AuNCs (excitation-450nm) in dispersed (black line) and in phase separated liquid condensate form (Red line).

Enhanced emission intensity after confinement of BSA@AuNCs within liquid droplets is observed***.*** The enhanced steady-state fluorescence emission in case of phase separated nanoclusters confirms the congregation of BSA@AuNCs within the liquid droplets.

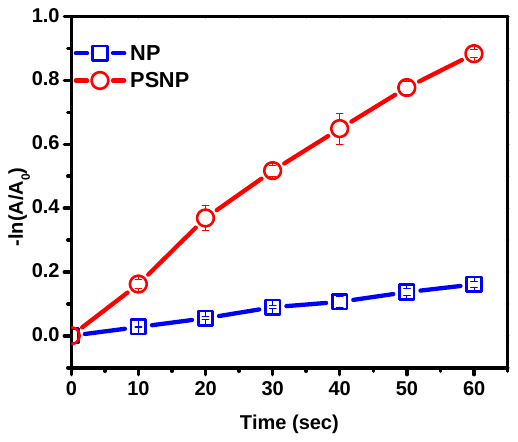

**Figure S5:** First order decay kinetic fit of reduction reaction of p-nitrophenol using BSA capped gold nanoparticles in dispersed (NP, blue line) and in phase separated condensate form (PSNP, red line).

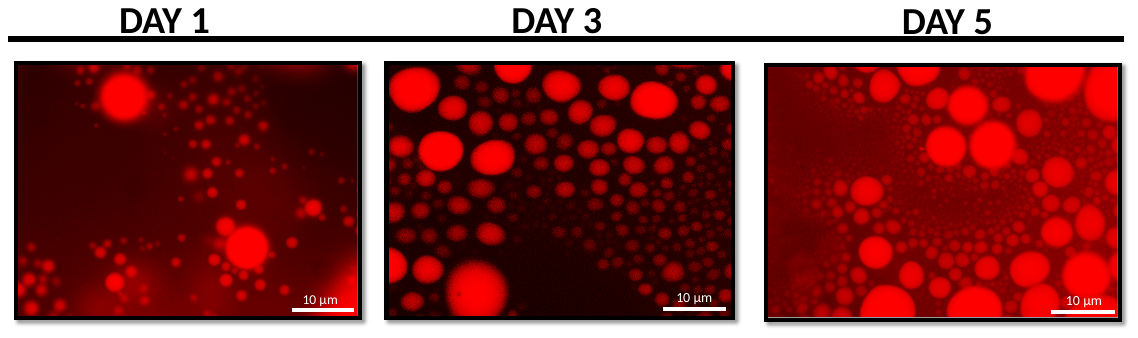

**Figure S6:** Day dependent visualisation of BSA@AuNCs liquid droplets

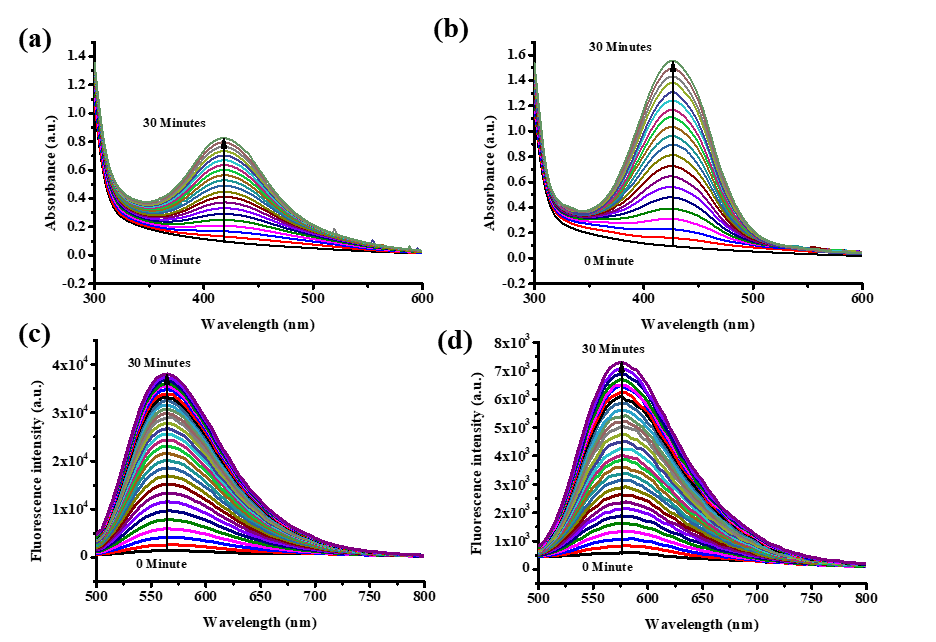

**Figure S7:** Increase in absorbance at 420nm with increase in oxidation of substrate o-phenylenediamine to product diaminophenazine (a) dispersed state of BSA@AuNCs, (b) separated state of BSA@AuNCs; increase in emission intensity at 565nm upon 420nm excitation (c) dispersed state (d) phase separated. This reveals the more product formation in case of phase separated BSA@AuNCs.

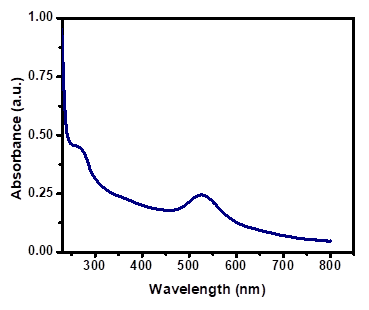
**Figure S8**: Absorbance spectra of α-synuclein coated citrate capped AuNPs.

.
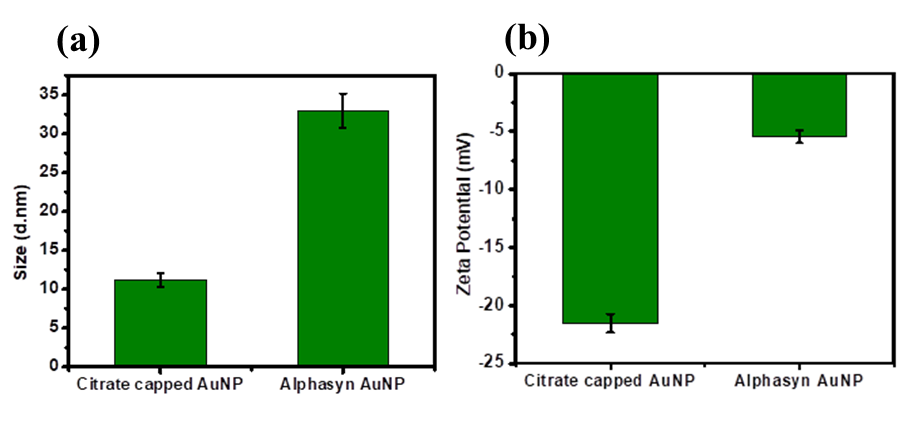
**Figure S9:** (a) Change in the hydrodynamic diameter of citrate capped AuNPs after conjugation with α-synuclein, (b) decrease in the zeta potential of citrate capped AuNPs due to interaction with positively charged N-terminal of α-synuclein protein.

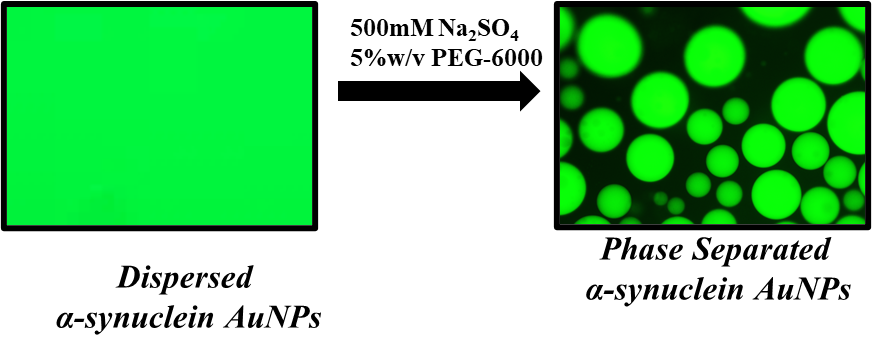

**Figure S10:** (a) Microscopic visualisation of liquid droplets of α-synuclein AuNPs in presence of 500mM Na_2_SO_4_ and 5%w/v PEG-6000.

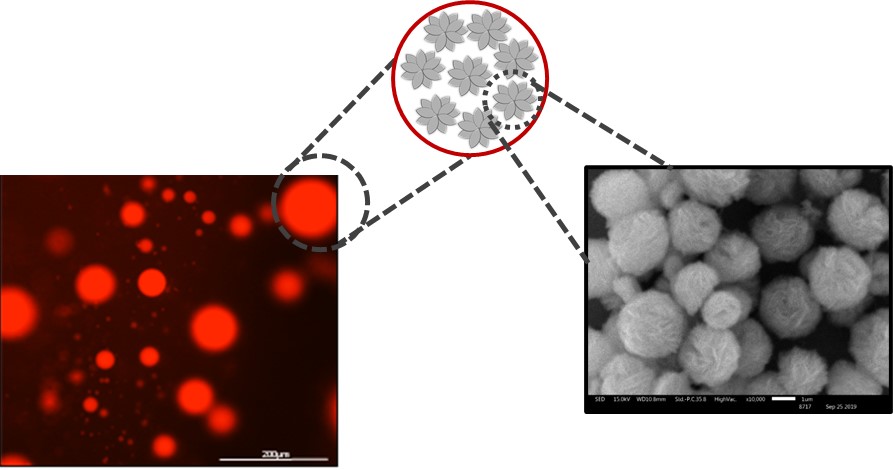

**Figure S11:** Liquid-liquid phase separation of BSA-CuPO_4_ nanoflowers in presence of 500mM sodium sulphate and PEG-6000 (5% w/v); SEM image of BSA-Copper phosphate nanoflowers showing its floral structure, Scale bar is-1µm

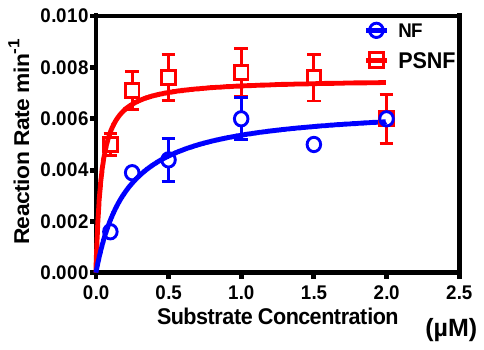

**Figure S12:** Oxidase activity of BSA-Copper phosphate nanoflowers for oxidation of pyrogallol to purpurogallin as dispersed solution (blue line) and in phase separated form (red line). Graph represents the Michaelis-Menten Fit for the kinetics data for 60-minute kinetic reaction.

| **Table S1: The values for K_m_ and V_max_ of BSA-Copper phosphate Nanoflower and Phase separated Nanoflowers** | | |
| --- | --- | --- |
|  | **K_m_ (µM)*** | **V_max_ (Ms^-1^)*** |
| **NF** | **0.22±0.01** | **0.06±0.001** |
| **PSNF** | **0.04±0.001** | **0.08±0.001** |
| A lower K_m_ in the case of phase separated nanoflowers indicate a higher affinity towards the substrate while, the comparable Vmax values in both the cases indicate the phase separation does not alter the enzyme kinetic rate. | | |
